# Distinct impacts of each anti-anti-sigma factor ortholog of the chlamydial Rsb partner switching mechanism on development in *Chlamydia trachomatis*

**DOI:** 10.1101/2024.02.22.581593

**Authors:** Shiomi Junker, Vandana Singh, Aamal G.M. Al-Saadi, Nicholas A. Wood, Scott D. Hamilton-Brehm, Scot P. Ouellette, Derek J. Fisher

## Abstract

Partner Switching Mechanisms (PSM) are signal transduction systems comprised of a sensor phosphatase (RsbU), an anti-sigma factor (RsbW, kinase), an anti-anti-sigma factor (RsbV, the RsbW substrate), and a target sigma factor. *Chlamydia* spp. are obligate intracellular bacterial pathogens of animals that undergo a developmental cycle transitioning between the infectious elementary body (EB) and replicative reticulate body (RB) within a host-cell derived vacuole (inclusion). Secondary differentiation events (RB to EB) are transcriptionally regulated, in part, by the house-keeping sigma factor (σ^66^) and two late-gene sigma factors (σ^54^ and σ^28^). Prior research supports that the PSM in *Chlamydia trachomatis* regulates availability of σ^66^. Pan-genome analysis revealed that PSM components are conserved across the phylum Chlamydiota, with *Chlamydia* spp. possessing an atypical arrangement of two anti-anti-sigma factors, RsbV1 and RsbV2. Bioinformatic analyses support RsbV2 as the homolog to the pan-genome conserved RsbV with RsbV1 as an outlier. This, combined with *in vitro* data, indicates that RsbV1 and RsbV2 are structurally and biochemically distinct. Reduced levels or overexpression of RsbV1/RsbV2 did not significantly impact *C. trachomatis* growth or development. In contrast, overexpression of a non-phosphorylatable RsbV2 S55A mutant, but not overexpression of an RsbV1 S56A mutant, resulted in a 3 log reduction in infectious EB production without reduction in genomic DNA (total bacteria) or inclusion size, suggesting a block in secondary differentiation. The block was corroborated by reduced production of σ^54/28^-regulated late proteins and via transmission electron microscopy.

**Importance:** *C. trachomatis* is the leading cause of reportable bacterial sexually transmitted infections (STIs) and causes the eye infection trachoma, a neglected tropical disease. Broad-spectrum antibiotics used for treatment can lead to microbiome dysbiosis and increased antibiotic resistance development in other bacteria, and treatment failure for chlamydial STIs is a recognized clinical problem. Here, we show that disruption of a partner switching mechanism (PSM) significantly reduces infectious progeny production via blockage of RB to EB differentiation. We also reveal a novel PSM expansion largely restricted to the species infecting animals, suggesting a role in pathogen evolution. Collectively, our results highlight the chlamydial PSM as a key regulator of development and as a potential target for the development of novel therapeutics to treat infections.

## Introduction

The phylum Chlamydiota is comprised of obligate intracellular Gram negative bacteria infecting a variety of eukaryotic hosts such as amoeba, invertebrates, and vertebrates including fish, reptiles, birds, and mammals (1). Like most obligate intracellular bacteria, they have reduced genomes varying in size from 1 Mbp to ∼2.5 Mbp, and genome content impacts their endosymbiotic host range by dictating, for example, metabolic capacity/parasitic requirements, cell entry mechanisms, stress response pathways, and protein effector repertoires needed to survive in the face of different innate and adaptive immune responses (2–5). Despite variation in hosts and endosymbiotic requirements, all characterized members of the phylum undergo a developmental cycle involving differentiation of the infectious elementary body (EB) into the replicative reticulate body (RB) (primary differentiation) followed by replication of RBs and the eventual conversion of RBs into EBs (secondary differentiation) (6, 7). Halting developmental progression would be an effective means of preventing or treating infections, but the mechanisms governing and executing differentiation are poorly understood.

Partner switching mechanisms (PSMs) are protein-phosphorylation based bacterial signal transduction systems that bacteria employ to sense and respond to environmental cues. <u>R</u>egulator of <u>S</u>igma <u>B</u>, or Rsb, is a well characterized PSM mostly found in Gram positive bacteria where it is used to sequester the availability of σ^B^ in response to environmental or energy stressors (8–13). Release of the regulated sigma factor allows for transcriptional responses that can lead to developmental changes including spore formation, activation of virulence genes, motility, biofilm formation, or initiation of various stress response pathways (14–22). Nominally, a PSM is composed of an anti-sigma factor with kinase activity (ASF; RsbW), a sensor phosphatase (RsbU), and an anti-anti-sigma factor (AASF; RsbV) that is the substrate for both RsbW and RsbU (23, 24). The ASF switches between a complex with the sigma factor (transcription repressed) and the unphosphorylated AASF (sigma factor freed, transcription competent) (25–28) (see Figure 1). Phosphorylation of the AASF by the ASF allows the ASF to switch back to the sigma factor (28–30). Activation of RsbU resets the AASF through dephosphorylation, allowing for complex formation with the ASF (31–33). In this way, transcription can be regulated in response to environmental cues or energy stress (34, 35).

**Figure 1.**
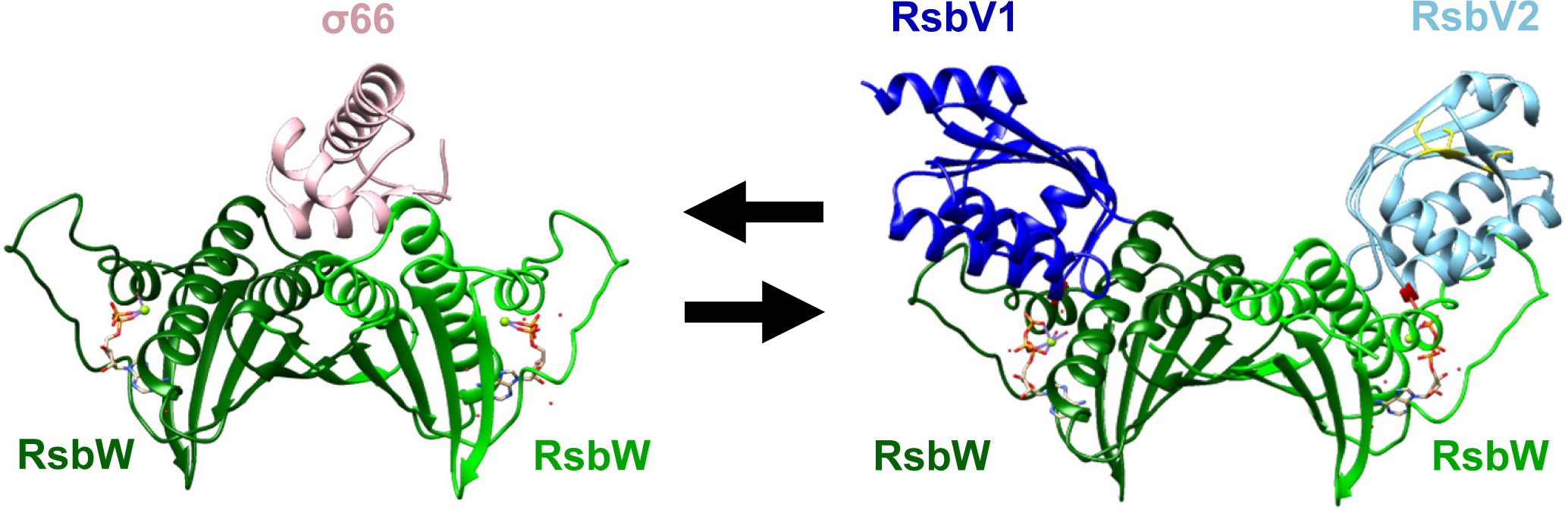
*In silico* structure models show RsbW switching protein interaction partners between σ^66^ and the AASFs RsbV1/RsbV2. Monomeric protein structures were predicted using AlphaFold. Each protein structure is superimposed with *B. subtilis* homologs SpoIIAB-σ^B^ binding with ADP, Mg^2+^, H_2_O (1L0O) or SpoIIAB-SpoIIAA with ATP, Mg^2+^, H_2_O (1TID), and the structures were rendered using UCSF chimera. RsbW (dark green) is dimerized with another RsbW (light green), which interacts with a part of σ^66^ (pink) or AASFs (RsbV1 as dark blue and RsbV2 as sky blue). An ATP molecule is present in each RsbW monomer. The phosphorylation site of each AASFs is highlighted in red (S56 for RsbV1, S55 for RsbV2). The RsbV2 CXCC motif is shown in yellow.

The phylum Chlamydiota genomes contain a core PSM, which includes RsbU, RsbW (ASF), and RsbV (AASF) (36). Studies of the PSM from *Chlamydia trachomatis* indicate that RsbU can sense the TCA-metabolite α-ketoglutarate and glycolysis intermediate phosphoenolpyruvate, that RsbW can phosphorylate RsbV in an ATP-dependent manner, that RsbU can dephosphorylate RsbV, and that RsbW can complex with σ^66^ (housekeeping sigma factor, σ^70^ family member) (37–41). In *C. trachomatis* and other *Chlamydia* that infect vertebrates, there are two additional sigma factors, σ^54^ and σ^28^, that promote transcription of late genes involved in secondary differentiation (42–46). The ratio of σ^66^-RsbW to RsbW-RsbV in the bacterium impacts transcription by altering the pool of RNAP core available to σ^28^ and σ^54^ and would have a titration effect on σ^66^-dependent gene transcription with priority given to unrepressed consensus promoters when free σ^66^ levels are reduced. In this manner, the chlamydial PSM can potentially regulate growth rate and development in response to nutritional cues.

Members of the genus *Chlamydia* are all pathogens of animals and include *C. trachomatis*, which is the leading cause of reportable bacterial sexually transmitted infections (47), and *Chlamydia pneumoniae*, which causes pneumonia and has been associated with increased risk of cardiovascular disease (48, 49). Broad spectrum antibiotics are used to treat chlamydial infections in humans and animals and can lead to dysbiosis of the microbiome (50). While antibiotic resistance is not currently a clinical problem, with the possible exception of tetracycline resistance observed in *Chlamydia suis* (pathogen of pigs), the bacteria can enter into a state of persistence leading to reduced sensitivity to currently used antibiotics that can result in treatment failure (51–54). Consequently, there is a need for improved therapeutic approaches for treating chlamydial infections.

Prior work from our lab and others’ has shown that genetic disruption of the PSM in *C. trachomatis* reduces the production of infectious progeny, suggesting that the PSM could represent a novel therapeutic target (38, 39, 41). Here, we report on variations of the PSM across the phylum Chlamydiota and hypothesize that AASF expansion and alterations helped facilitate speciation of the animal-infecting *Chlamydia* (“pathogenic *Chlamydia*”). We also provide additional evidence supporting the hypothesis that the PSM functions in regulating secondary differentiation and that genetic manipulation of the PSM cripples production of EBs.

## Results

### AASF gene copy number (RsbV versus RsbV1 and RsbV2) varies across the Chlamydiota phylum

*C. trachomatis* and the other members of the genus *Chlamydia* (all pathogens of animals) possess two copies of RsbV, denoted RsbV1 and RsbV2, and a putative second sensor phosphatase, dubbed CTL0852 in *C. trachomatis* L2. Possession of dual AASFs is atypical, and is especially intriguing for a minimal genome organism such as *Chlamydia* (∼1 to 1.3 Mbp) (3). Our group and others have shown that both RsbV1 and RsbV2 can be phosphorylated by RsbW on conserved serine residues, S56 and S55, respectively, and that both AASFs can be dephosphorylated by RsbU (37, 38, 40, 41). However, the kinetics of phosphorylation and dephosphorylation towards RsbV1 are more favorable *in vitro* compared to RsbV2 and a mechanistic explanation for those differences is lacking.

Modeling of the interactions between RsbW-σ^66^ and RsbW-RsbV1 or RsbW-RsbV2 reveals that both AASFs should be able to disrupt σ^66^-RsbW interactions (Figure 1). While the phosphorylation site motifs and AASF/ASF interactions sites in RsbV1 and RsbV2 are mostly conserved, the models indicate differences in a number of amino acids at the RsbW-AASF interface that would alter the charge profile, potentially impacting RsbW-AASF binding and/or phosphorylation (Figures S1 and S2). RsbV1 also has an additional six C-terminal amino acids (116 amino acids) versus RsbV2 (110 amino acids), and the proteins have predicted pI’s that vary by two logs (pI of 5 for RsbV1 [positively charged in the chlamydial cytoplasm] and a pI of 7 for RsbV2 [no net charge], (55)). We attempted to directly assess the importance of amino acids predicted to be involved in RsbW-AASF binding using amino acid point mutants, but the recombinant proteins were either insoluble or did not properly fold as measured by limited trypsin proteolysis (data not shown). As an alternative approach, time course kinase and phosphatase assays were performed at different pH values. Similar trends were observed as previously observed (37, 38, 40, 41), with RsbV1 functioning as a preferred kinase (Figure S3A) and phosphatase substrate (Figure S3B).

While *Chlamydia* possesses RsbV1 and RsbV2, the presence of dual AASFs is not universal across Chlamydiota, and most phylum members only have a single AASF (5). Given the differences in RsbW/RsbU affinities towards RsbV1/RsbV2 and the divergence in genome conservation, we sought to identify which AASF represented the “ancestral” copy. Phylogenetic (Figure 2A) and CLANS analyses (Figure 2B) along with gene synteny (Figure 2C) suggest that the RsbV2 in *Chlamydia* is more similar to the single AASF, RsbV, found in non-*Chlamydia* members. This would make RsbV1 unique to the chlamydial organisms most commonly associated with infections in animals. In addition, the amino acid sequence identity of RsbV2/RsbV across species shows more variability than when comparing RsbV1 across species (Figure S1 and Table S1). We speculate (i) that the RsbV2/V variability is linked to the longer evolutionary possession of this AASF and the diverse environments/hosts associated with these bacteria and (ii) that RsbV1 serves as an “expansion factor” involved in the leap to becoming an animal pathogen. For example, comparing *C. trachomatis* and *C. pneumoniae,* RsbV1 exhibits 71% similarity while RsbV2 has only 44% similarity. Relevant to our proposed dichotomy regarding the number of AASFs between pathogenic and “non-pathogenic” *Chlamydia*, there are some discrepancies in the literature and annotated genomes (5). While the following pathogenic chlamydia have been annotated as possessing one AASF, our reassessment of the deposited genome sequences supports the presence of two AASFs for *C. abortus*, *C. pecorum*, *C. ibidis*, and *Candidatus* Clavichlamydia salmonicola (Table S2). Additionally, there are two “non-pathogenic” species annotated to have two copies of AASFs from the Amoebachlamydiae, *Parachlamydiaceae* bacterium HS-T3 and *Parachlamydia* sp. C2. The phylogenic tree (Figure 2A) shows that both of these AASFs are more similar to RsbV2 than RsbV1.

**Figure 2.**
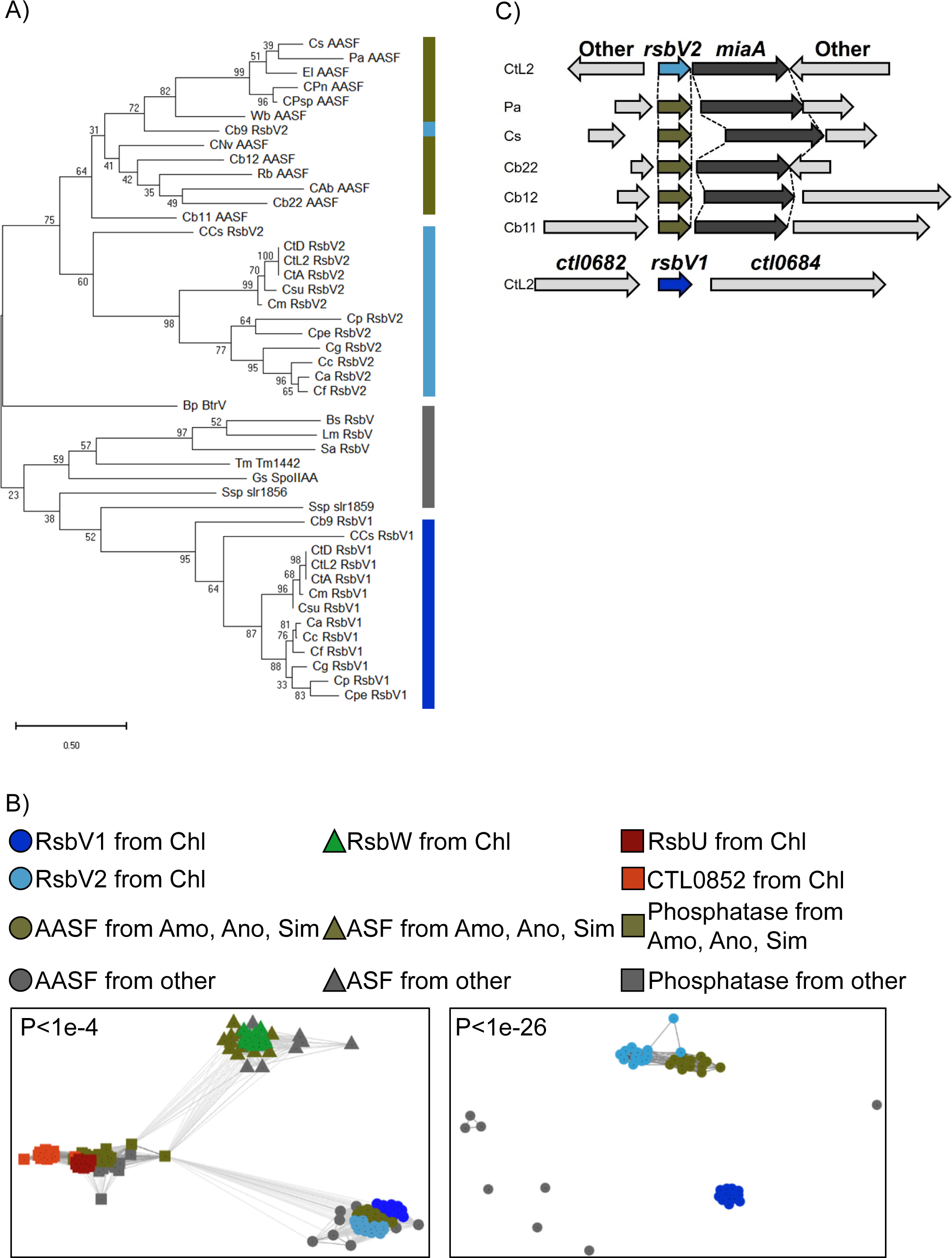
*In silico* studies reveal a potential AASF expansion. PSM homologs (RsbV, RsbW, and RsbU) were obtained from Chlamydiales (Chl), Amoebachlamydiales (Amo), Anoxychlamydiales (Ano), Simkaniales (Sim), and non-chlamydiia species (Other) for *in silico* analyses. NCBI IDs of each protein and species abbreviations are listed in Table S2. A) A phylogenetic tree of AASFs was constructed in MEGA11 using the Maximum likelihood method with 1000 bootstraps. Numbers represent the percentage of trees in which the associated taxa clustered together. B) Protein sequence relationships were also analyzed in CLANS. The left panel describes PSM proteins with the connection defined as P < 1e^-4^, while the right panel shows AASF connections only using a P < 1e^-26^. C) Gene synteny analysis of *rsbV2* (*rsbV*). Gene loci were obtained from the NCBI genome database (see Table S2 for NCBI IDs).

### Phosphorylation status of the AASFs throughout development

We previously reported that RsbV1 and RsbV2 protein levels do not appear to shift relative to MOMP under different glucose levels and that RsbV2 levels do not change in an RsbV1-null mutant (41). Soules et al. found that an RsbU-null strain (PSM phosphatase deficient) was severely attenuated for growth and IFU production compared to wild type and *rsbU*-complemented strains (39). In addition, Fisher et al. detected phosphorylated RsbV1 and RsbV2 in EBs, but not in RBs, for *C. caviae* (56). Collectively, these data support phosphorylation of the AASFs as an important driver of PSM regulation, leading us to investigate phosphorylation status during development. Our previous *in vitro* work with RsbV1 and RsbV2 used His-tagged constructs, but we were unable to detect His-tagged AASFs expressed in *C. trachomatis* by western blot necessitating a switch to a 3XFLAG tag. We initially explored the phosphorylation status of RsbV2 during infection using a recombinant *C. trachomatis* strain overexpressing a C-terminal FLAG-tagged RsbV2 (RsbV2-FLAG). Samples were separated on Phos-tag acrylamide gels to resolve phosphorylated and non-phosphorylated proteins, and RsbV2 was detected via anti-FLAG western blot. Only phosphorylated RsbV2-FLAG was detected even at the 24 hpi sample when RB levels should be abundant (Figure S4A). *In vitro* kinase and phosphatase assays with N-terminal and C-terminal FLAG-tagged RsbV2 showed that phosphatase activity was impaired compared to activity towards His-P-RsbV2 (Figure S4B and C). Attempts to detect endogenous RsbV2 using Phos-tag acrylamide gels and anti-RsbV2 western blot were only successful with samples harvested at 48 hpi (EB-enriched), and detection required RsbV2 immunoprecipitation from bacteria-enriched samples. Similar to the *C. caviae* EB samples (56), endogenous P-RsbV2 was detected from EB-enriched *C. trachomatis* infected cells (Figure S4D). Using a similar approach as for FLAG-RsbV2, N-terminal FLAG-RsbV1 appeared to be phosphorylated at all time points beyond 15 hpi (earliest time at which protein could be detected) (Figure S4E). Attempts to detect endogenous RsbV1 using Phos-tag acrylamide gels, bacteria-enriched samples, and anti-RsbV1 western blot were unsuccessful.

### RsbV2 knockdown or overexpression does not impact EB production

Previous work with an RsbV1-null strain (*rsbV1*::GII intron) showed that IFU production dropped by ∼1 log compared to the parental strain (38, 41). Attempts to generate an *rsbV2* null mutant have been unsuccessful, so we utilized CRISPRi-mediated dCas12 knockdown of *rsbV2* as an alternative approach (57). We also overexpressed FLAG-tagged RsbV2 to assess the impact of increased levels of RsbV2 on chlamydial development. Knockdown of *rsbV2* in RsbV1-competent or RsbV1-null strains did not lead to a significant reduction in IFU production at 24 or 48 hpi compared to non-targeting and uninduced controls (Figure 3A). A downward trend in infectious EB production was observed compared to uninduced and non-targeting samples at 48 hpi in both RsbV1-competent and RsbV1-null strains (Figure 3A). RsbV2 western blots confirmed knockdown of *rsbV2* (Figure 3B). Since significant changes were not observed in inclusion morphology (data not shown) or in IFU production, we did not measure changes in *miaA* levels (putative operon with *rsbV2*, Figure 2B) or perform complementation experiments. Overexpression of FLAG-RsbV2 under high glucose or low glucose conditions also resulted in no significant changes in IFU production (Figure 3C). Consistent with prior results, overexpression of RsbV1 (as FLAG-RsbV1 in this study) resulted in an upward trend in IFU production. Growth under different glucose conditions was tested as the PSM may function to set a growth cap and/or govern a developmental timeline in response to the levels of metabolites such as ATP, α-ketoglutarate, and PEP (37–41).

**Figure 3.**
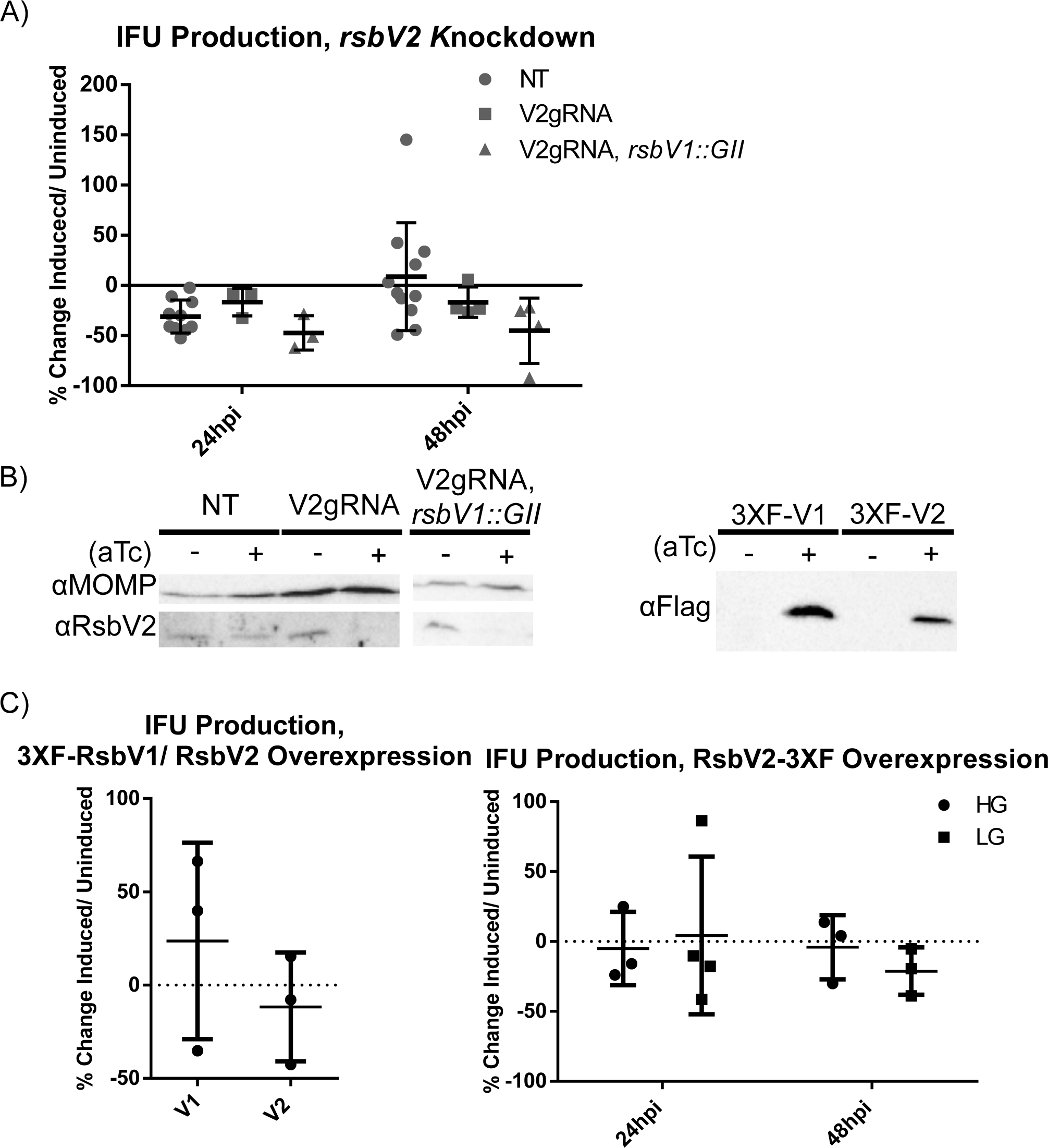
RsbV2 overexpression or *rsbV2* knockdown does not alter IFU production. Recombinant *C. trachomatis* L2 (wild type chromosome or an *rsbV1*-inactivated strain [*rsbV1*::GII]) with CRISPRi dCas12 non-targeted (NT) or *rsbV2*-targeted (V2gRNA) plasmids were used to infect HeLa cells. Knockdown of *rsbV2* was induced by 2 nM aTc at 4 hpi, and samples were harvested at 24 or 48 hpi for infectious EB determination by IFU assay (A). Bars represent the mean, and the error bars report standard deviation. Statistical analysis was done by two-way ANOVA, there is no significant difference. B) Western blotting was performed to confirm RsbV2 knockdown (left panel) or RsbV2/RsbV1 overexpression (right panel). MOMP was used as a western blot loading control. Primary antibodies used are indicated to the left of each blot. C) N-ter 3X FLAG-tagged RsbV1 or RsbV2 were induced with 10 nM aTc in high glucose medium, and samples were harvested at 48 hpi for titering (left panel). C-ter 3X FLAG-tagged RsbV2 was induced with 10 nM aTc at 4 hpi in infected cells cultured in either high glucose or low glucose, and samples were harvested at 24 hpi or 48 hpi for titering by IFU assay (right panel). Bars represent the mean, and error bars report the standard deviation. Statistical analysis was done by t-test, and there is no significant difference. IFUs were normalized to uninduced samples for (A) and (C).

### Expression of a non-phosphorylatable RsbV2 S55A mutant significantly reduces IFU production but not bacterial numbers

To assess the importance of reversible AASF-RsbW interactions, we expressed non-phosphorylatable RsbV1 (S56A) and RsbV2 (S55A) analogs. While detection of only phosphorylated RsbV1 and RsbV2 during infection was unexpected (Figure S4), and subject to experimental limitations, our results do confirm that RsbW is active during infection and that the FLAG-tag does not prevent RsbW-AASF interactions and phosphorylation *in vivo*. Previous data from surface-plasmon resonance experiments (38) and yeast two-hybrid (37) or bacterial adenylate cyclase two-hybrid experiments (data not shown, (58)) demonstrated that RsbW forms a more stable complex with the RsbV1 S56A and RsbV2 S55A mutants than with the wild type proteins. “Trapping” of RsbW by an AASF should increase the pool of free σ^66^. In our model, this leads to competition with late-gene sigma factors (σ^54/28^) for binding with the RNAP core and/or alterations in transcription of σ^66^-dependent genes via promoter titration. Competition and promoter titration would be predicted to interfere with growth and/or secondary differentiation.

Expression of an N-terminal FLAG-RsbV1 S56A mutant in a wild type background strain resulted in no significant differences in IFU production at 24 or 48 hpi (Figure 4). Unexpectedly, FLAG-RsbV1 S56A expression in the RsbV1-null strain resulted in a significant increase in IFU production of ∼2.5-fold at both time points. To assess if the FLAG-tag location impacted the results for RsbV1 S56A, we also tested a C-terminal FLAG-tagged RsbV1 S56A. For reasons that we cannot explain, expression of the C-terminal tagged construct was not detectable via anti-FLAG western blot even with increased aTc-inducer levels (112 nM vs 10 nM used in other experiments). RsbV1-FLAG S56A expression could be detected via immunofluorescence, although the amount of RsbV1-FLAG S56A produced was well below the levels of RbsV2-FLAG S55A expressed under equivalent induction conditions (Figure S5). Expression of the RsbV1-FLAG S56A did result in a 50% reduction in IFUs compared to the uninduced strain. However, the reduction was not significantly different when compared to a GFP-FLAG expressing strain used as a control for the high amount of aTc used for induction (Figure 4B). Since expression of the RsbV1 S56A in the RsbV1-null strain had a modest rescuing effect on IFU production comparing wild type and RsbV1-null IFU production levels, we expressed RsbV2-FLAG S55A in the RsbV1-null background. In contrast to the RsbV1 S56A results (rescue), IFU production was decreased ∼3 log when expressing the RsbV2-FLAG S55A mutant (Figure 4C), and similar results were obtained upon expression of RsbV2-FLAG S55A in a wild type background (RsbV1 competent). The reduction in IFUs when expressing the RsbV2-FLAG S55A mutant in the RsbV1-null strain was greater than when expressing RsbV2-FLAG S55A in the wild type background (RsbV1/RsbV2 competent strain). The severe IFU reduction resulting from RsbV2 S55A overexpression led us to further define the phenotype.

**Figure 4.**
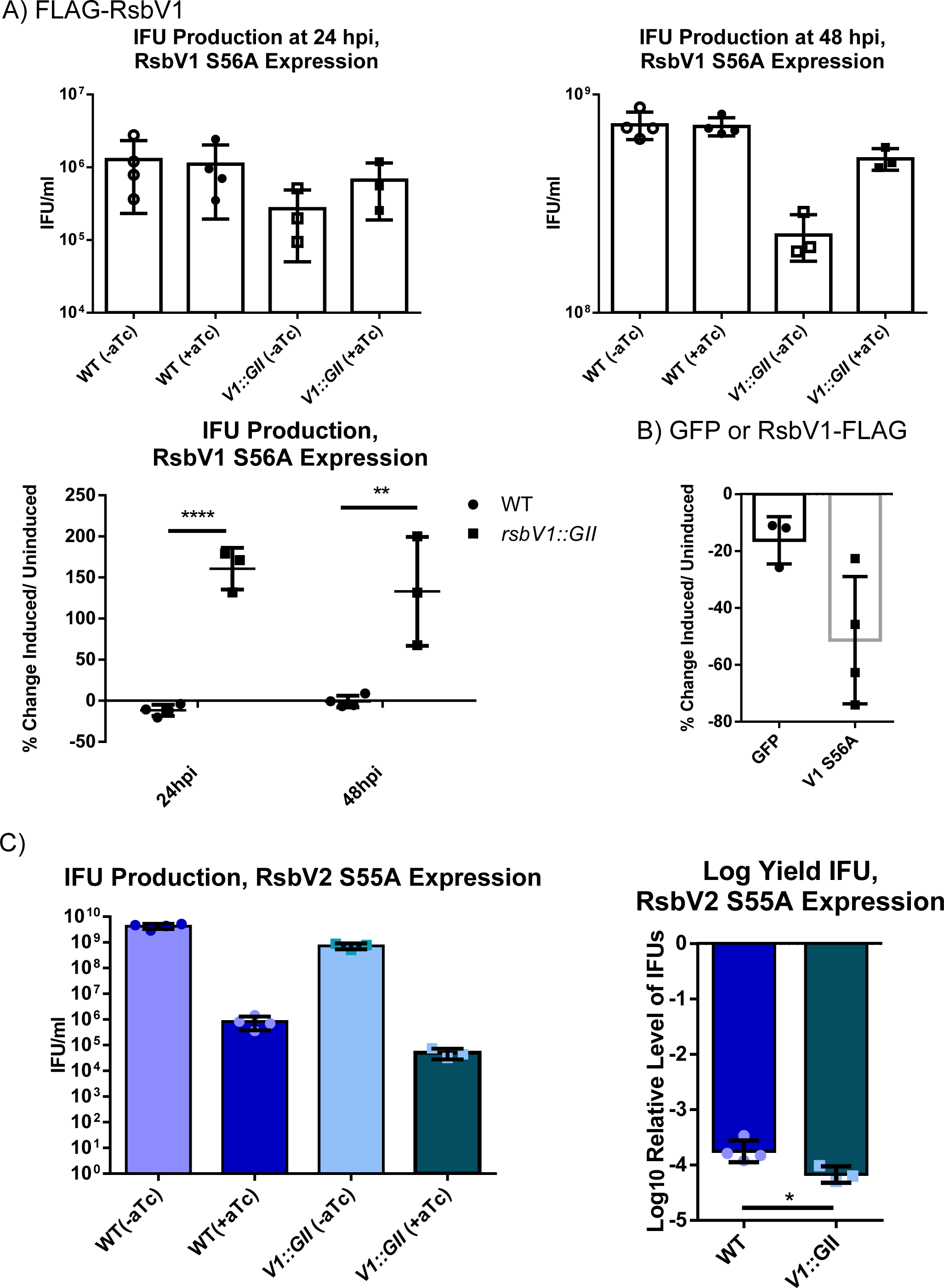
Induction of a non-phosphorylatable RsbV1 S56A mutant does not alter IFU production. RsbV1 S56A or RsbV2 S55A were induced with aTc at 4 hpi in recombinant *C. trachomatis* L2 (wild type chromosome or an *rsbV1*-inactivated strain [*rsbV1*::GII]). Titers were assessed for infectious progeny numbers at 24 or 48 hpi using the IFU assay. A) N-ter 3X FLAG-tagged RsbV1 S56A induced with 10 nM aTc, B) C-ter 3X FLAG-tagged GFP (control) or RsbV1 S56A induced with 112 nM aTc. Statistical analysis shows no significant difference. C) C-ter 3X FLAG-tagged RsbV2 S56A was induced with 4 nM aTc in a wild type chromosome background or an *rsbV1*-inactivated strain [*rsbV1*::GII] at 4 hpi, and harvested at 48 hpi. Statistical analysis was done with the t-test (ns = P > 0.05, * = P ≤ 0.05, ** = P ≤ 0.01, **** = P ≤ 0.0001).

Expression of an RsbV2-FLAG S55A mutant resulted in a significant drop in IFU production with an approximate 3 log reduction observed at 36 hpi and beyond compared to the uninduced sample or a strain overexpressing wild type RsbV2-FLAG (Figure 5A and B). Allowing infections to proceed for 72 hpi did not lead to recovery of IFU numbers in the presence of RsbV2-FLAG S55A. In contrast to the significant reduction in IFUs, bacterial numbers were not significantly different between RsbV2 S55A induced and uninduced samples based on quantification of genomic DNA (Figure 5C and D). Inclusion sizes measured using immunofluorescence microscopy were not grossly altered when expressing RsbV2-FLAG S55A, with a modest size increase found at 48 hpi (Figure 5E and Figure S6). Experiments performed with an N-terminal His-tagged RsbV2 S55A mutant phenocopied the IFU results for the RsbV2-FLAG S55A mutant (Figure S7). The impact of RsbV2 S55A expression on growth under low glucose conditions could not be measured owing to the already reduced IFU production of the uninduced strain during low glucose conditions (data not shown).

**Figure 5.**
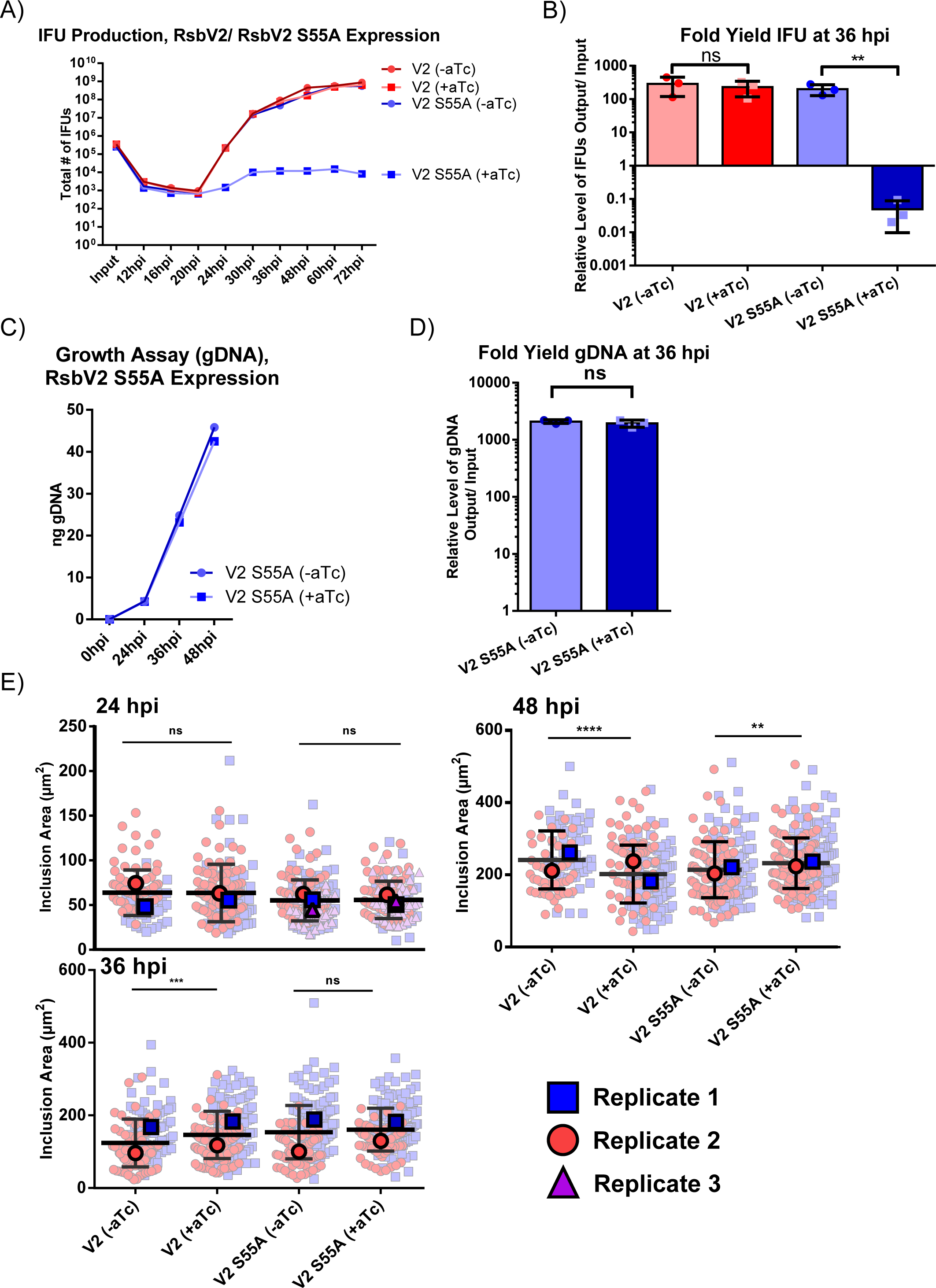
Expression of a non-phosphorylatable RsbV2 S55A mutant significantly reduces IFU production. Recombinant *C. trachomatis* L2 with an aTc-inducible C-ter 3X FLAG-tagged wild type or S55A RsbV2 were used to infect HeLa cells. RsbV2 was induced by adding 10 nM aTc at 4 hpi. Samples were harvested at different time points and used for western blotting and assessment of IFU production. A) Titers of infectious EBs were determined using an IFU assay and fold yield for the 36 hpi time point is reported in (B). Note that the IFU fold yield for the induced RsbV2 S55A mutant strain was statistically different at 36-72 hpi, while the induced wild type RsbV2 strain was not. C) The number of bacteria (RBs+EBs) were quantified by qPCR and fold yield of gDNA at 36 hpi is represented in D. Plots in (A) and (C) are representative of three experiments, (B) and (D) report the averages for all three trials. E) Immunofluorescence microscopy was performed with anti-MOMP antibody and an anti-mouse secondary conjugated with Alexa Fluor 488 to demarcate inclusions. Images were acquired at 1000x magnification and the size of inclusions was measured using ImageJ. Pale symbols report individual inclusion size, dark symbol represents the average of each replicate. Statistical analyses were done with the t-test (ns = P > 0.05, ** = P ≤ 0.01, *** = P ≤ 0.001, **** = P ≤ 0.0001).

### Expression of RsbV2 S55A blocks chlamydial developmental

The reduction in IFUs without gross alterations in inclusion morphology or reduced levels of genomic DNA indicated that chlamydial development was being perturbed with either lack of EB production and/or the production of non-infectious EBs. We initially used western blot to determine if production of late-stage proteins had been altered (Figure 6A). We assessed three late proteins regulated by the three different sigma factors: the histone-like proteins HctA (regulated by σ^66^) and HctB (regulated by σ^28^) and the type 3 secretion effector Tarp (regulated indirectly by σ^54^) (42, 45, 46, 59). Protein levels were compared to MOMP as a marker for total bacteria (Figure 6B). For HctA, there was no difference when comparing uninduced and induced samples. However, levels of both HctB and Tarp were significantly reduced under induced conditions. These results are consistent with reduced amounts of RNAP-σ^54/28^ holoenzyme complexes or reduced levels of σ^54^/^28^ altogether and suggest that EB production is altered or blocked in the presence of RsbV2 S55A. We next used transmission electron microscopy to visualize the bacteria within the inclusion. Samples were imaged at 48 hpi and inclusions from induced samples showed relatively few EBs, numerous IBs and atypical IBs, and numerous RBs whereas uninduced samples contained mostly EBs (Figure 6C and Figure S8). In whole, our data support that expression of RsbV2 S55A leads to disruption of secondary differentiation with bacteria appearing to either remain as RBs or stall at the IB stage, which is representative of early secondary differentiation events. Extrapolating the protein production data for HctB and Tarp to other late proteins, we predict that secondary differentiation cannot be completed owing to reductions in σ^54/28^-regulated gene products.

**Figure 6.**
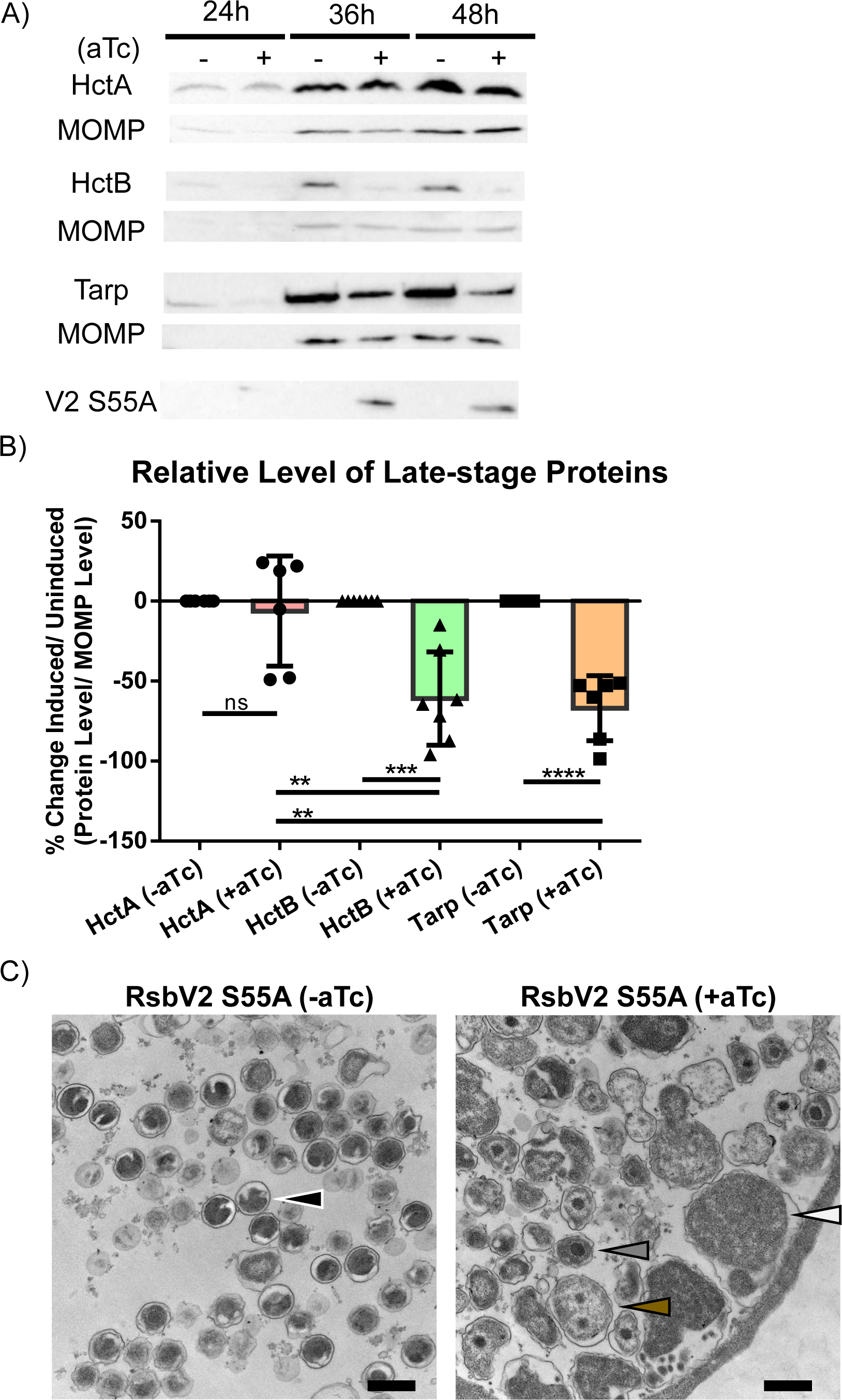
Induction of the RsbV2 S55A mutant reduces production of late-stage proteins and reduces EB formation. Recombinant *C. trachomatis* L2 producing C-ter 3X FLAG-tagged RsbV2 S55A was used to infect HeLa cells. RsbV2 was induced by adding 10 nM aTc and lysates were harvested at 24 hpi, 36 hpi, 48 hpi for western blot or samples were fixed at 48 hpi for TEM. A) Protein production was determined using quantitative western blot of induced or uninduced RsbV2 S55A samples. MOMP was used as a loading control. Proteins detected are listed to the left of each blot. B) Protein levels were quantified in reference to MOMP for HctA, HctB, and Tarp. Statistical analysis was done with the t-test by using HctA as a reference. (ns = P > 0.05, ** = P ≤ 0.01, *** = P ≤ 0.001, **** = P ≤ 0.0001) C) Uninduced (left image) or induced (right image) sample images were taken by TEM with magnification at 26,000x. Arrows denote a representative normal EB (black), IB (gray), abnormal IB (brown), and an RB (white). Additional TEM images are shown in Figure S8.

## Discussion

During chlamydial evolution, genome reduction has been balanced with gene acquisition allowing for select chlamydial species to gain the ability to infect vertebrates (5). One such expansion appears to have occurred within the chlamydial Rsb-like partner switching mechanism whereby the pathogenic *Chlamydia* possess two AASF homologs, RsbV1 and RsbV2. Here we sought to address why *Chlamydia* have two AASFs with different apparent affinities and reaction kinetics with the sensor phosphatase RsbU and the ASF, RsbW. We found that the majority of the phylum Chlamydiota have a single AASF whereas the Chlamydiales members possess two AASFs. CLANS, phylogeny, gene synteny, and sequence comparisons support that RsbV2 is the “ancestral” RsbV whereas RsbV1 is the acquired AASF. Note that we cannot rule out that both copies existed in a last universal common ancestor with RsbV1 being deleted in non-Chlamydiaceae members. It is also unclear as to where *rsbV1* would have originated given the somewhat “sheltered” life for obligate intracellular bacteria. Duplication and divergence would provide a simple mechanism, however such an event might be expected to result in genes in close proximity on the chromosome. There is synteny surrounding RsbV1 in positive-members, so multiple genes may have been acquired along with *rsbV1*. We hypothesize that dual AASFs played a role in the leap to infecting vertebrates, and both are found in the fish pathogen *Candidatus* Clavichlamydia salmonicola (along with the late gene regulator σ^28^ and various T3S effector proteins), a non-Chlamydiaceae member that has been proposed to be a transitional strain between environmental and pathogenic *Chlamydia* (4).

We remain left with the question as to what benefit a second AASF provides and how this altered PSM network facilities bacterial infection in vertebrates. RsbV1 and RsbV2 have different biochemical properties with disparate pI’s (5 and 7, respectively) and altered amino acids at the predicted AASF/ASF interaction site. We also note that RsbV2 possesses a CXCC motif (Figures 1 and S1, shown in yellow) that is absent in RsbV1 and the RsbV in non-Chlamydiaceae spp. During secondary differentiation, EBs become oxidized, which leads to cross-linking of numerous outer membrane proteins, and this has been speculated to impart stability to the infectious EB particle (60–62). It seems reasonable that redox changes are not localized to surface proteins and that they occur throughout the bacterium. The oxidoreductive state of numerous proteins would then serve as a regulatory mechanism controlling protein activity between the RB and EB forms. In the case of RsbV2, we predict that oxidation in the EB could prevent interactions with RsbW. RsbW could then sequester σ^66^, and we are exploring this hypothesis using *in vitro* and *in vivo* approaches.

Both RsbV1 and RsbV2 could impact PSM output via altered levels of the proteins themselves and/or through the pool of phosphorylated/nonphosphorylated RsbV1/RsbV2. Our prior work and data from the current study suggest that overall protein levels have a minimal impact on chlamydial growth and development. Inactivation of RsbV1 or knockdown of RsbV2 shows either a ∼1 log reduction in EB production or no significant reduction in EB production, respectively (38, 41). It is not clear if the different phenotypes observed are due to differential impacts of RsbV1 and RsbV2 within the PSM or from the different approaches whereby *rsbV2* knockdown is not equivalent to a gene knockout. Attempts to knockout *rsbV2* have been unsuccessful and could be attributed to technical problems, gene essentiality, or operon disruption as *rsbV2* is 5’ to *miaA* (a tRNA dimethylallyltransferase, not essential in other bacteria (63)). Unexpectedly, knockdown of *rsbV2* in an RsbV1-null background did not exacerbate the RsbV1-null phenotype. There is evidence for differential impacts of AASF deletion on bacterial phenotypes. Like the pathogenic *Chlamydia*, the cyanobacterium *Synechocystis* sp. PCC 6803 has two AASFs with Slr1856 a better *in vitro* RsbW substrate than Slr1859, and Slr1859 is not dephosphorylated *in vitro* by the RsbU homolog IcfG (64). Deletion of the “inferior” AASF, Slr1859, results in attenuated growth in inorganic carbon-limited medium supplemented with glucose whereas the Slr1856 deletion strain does not impact growth (65). Overexpression of either RsbV1 or RsbV2 also fails to elicit significant phenotypic changes, and we previously showed that growth under low glucose conditions to introduce nutritional stress did not lead to alterations in RsbV1/V2 levels. Our results are consistent with other bacteria in which perturbation of the AASF level had a minimal impact on the PSM-linked phenotype compared to alterations in the sensor phosphatase or ASF (34). Indeed, an RsbU-null *C. trachomatis* strain is highly attenuated for growth in cell culture (39). Collectively, the data support that phosphorylation status is the major driver of the AASF contribution to PSM regulation.

To explore the impact of AASF phosphorylation on growth and development, we first attempted to track RsbV1 and RsbV2 phosphorylation status over time using Phos-tag acrylamide and western blot to resolve and detect phosphorylated and non-phosphorylated species. Detection of phosphorylated RsbV1/RsbV2 from infected cells proved difficult likely owing to both protein abundance and technical problems revolving around differential protein transfer rates of phosphorylated and non-phosphorylated proteins when performing western blots with Phos-tag acrylamide gels. To improve sensitivity, we developed a new bacteria-enriching extraction method that should be useful for detecting other low abundant proteins. While we could detect phosphorylated FLAG-tagged RsbV2 starting at 24 hpi, we determined that the FLAG-tag interfered with RsbU-mediated dephosphorylation. Wild type RsbV2 could only be detected at 48 hpi, in which it appeared to be phosphorylated. RsbV1 was detected in the phosphorylated form and only the FLAG-tagged version could be detected. The presence of phosphorylated RsbV2 and RsbV1 in late infection samples reflects prior work from our group that only identified phosphorylated RsbV1 and RsbV2 in the EB form of *C. caviae* (56). Phosphorylated RsbV1 and RsbV2 is predicted to facilitate RsbW-σ^66^ interactions inhibiting transcription in EBs.

To perturb phosphorylation-mediated regulation of the AASFs, we overexpressed non-phosphorylatable forms of RsbV1 (S56A) and RsbV2 (S55A). While we did not observe significant phenotypes when overexpressing RsbV1 S56A, overexpression of RsbV2 S55A resulted in ∼3 log reduction in IFUs. Production levels of RsbV1 S56A were less than that of RsbV2 S55A even though the same expression vector was used (greater decrease observed for RsbV1-FLAG than for FLAG-RsbV1). We have observed this phenotype (differential expression levels from the same vector) for a number of genes and even when expressing amino acid point mutants (unpublished observation) suggesting potential differences in RNA stability and/or protein stability that occurs in a gene-dependent manner. We were initially surprised to observe a ∼3 log reduction in IFUs for the RsbV2 S55A overexpression strain as inclusion size was not obviously impacted. Quantification of gDNA under induced and uninduced conditions suggested that total numbers of bacteria were also similar in the presence or absence of RsbV2 S55A. Our model predicts that RsbV2 S55A should bind and sequester RsbW, freeing σ^66^ and leading to dysregulation of secondary differentiation. Potential mechanisms underlying dysregulation include a reduction in late gene expression through alterations in the levels of σ^54^ and/or σ^28^ or reduction in σ^54^/σ^28^-RNAP complexes due to competition with σ^66^ (Figure 7). Knockdown of σ^28^ or/and σ^54^ has been shown to be detrimental for IFU production, which phenocopies our results when overexpressing RsbV2 S55A (46). Over expression of other mutant proteins, such as a ClpX R230A mutant that cannot recognize trans-translation tagged proteins, has also been shown to block secondary differentiation suggesting that multiple mechanisms underlie RB to EB conversion (66).

**Figure 7.**
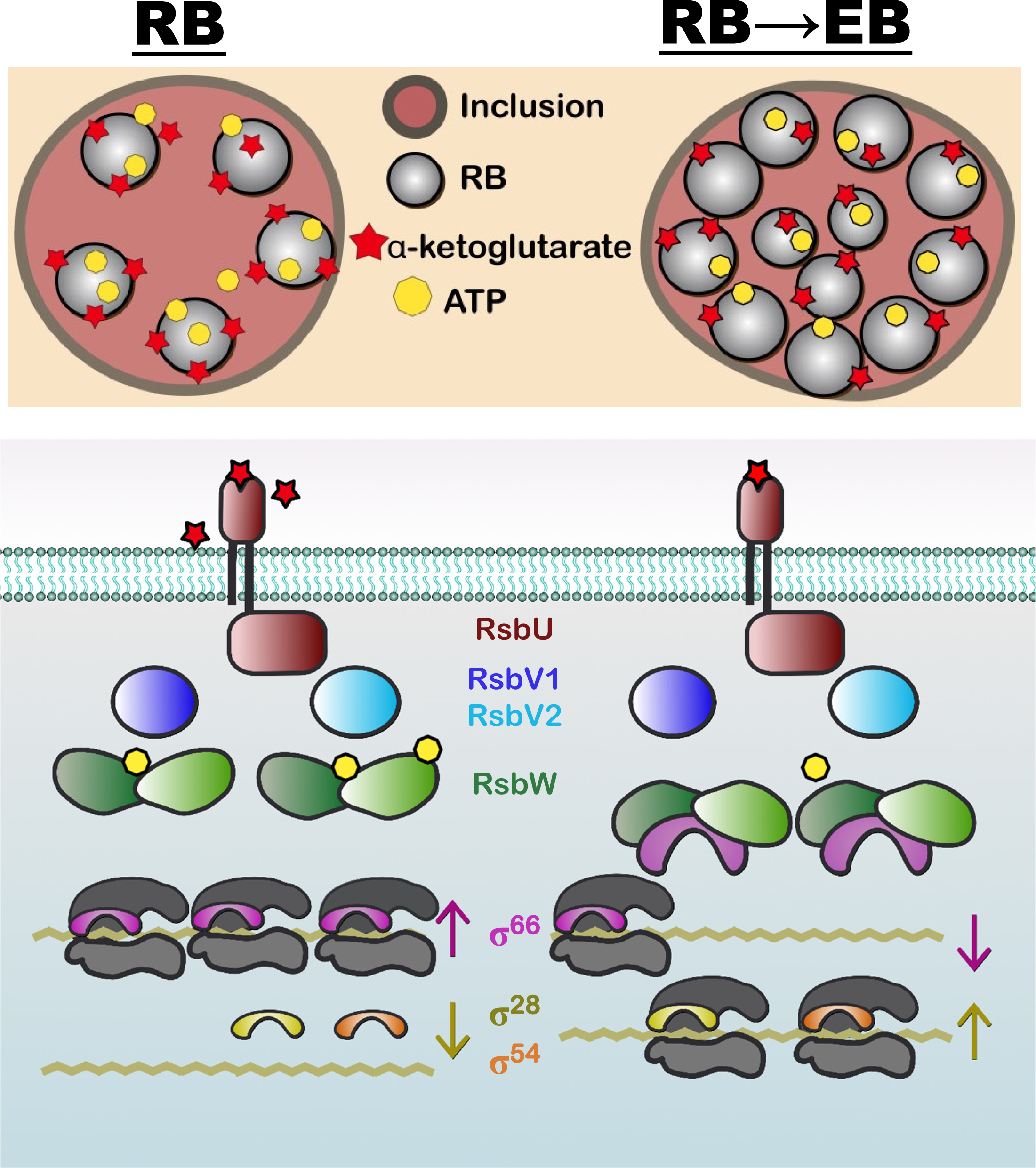
A working model of PSM-mediated developmental regulation. RsbW interacts with σ^66^ and represses transcription, or it phosphorylates RsbV1 and RsbV2 leading to derepression in an ATP-dependent manner. RsbU dephosphorylates both RsbV1 and RsbV2 in response to cell-derived, periplasmic α-ketoglutarate (prediction). During growth and replication, bacteria compete for nutrients including cell-derived ATP, glucose-6-P (can be used for substrate-level phosphorylation), and α-ketoglutarate. As bacterial numbers increase, the reduction of ATP and α-ketoglutarate levels decrease the signal to RsbU and RsbW leading to RsbW sequestering a pool of σ^66^, allowing alternative sigma factors (σ^28^ and σ^54^) to access the RNAP core supporting RB to EB conversion.

Consistent with a block in secondary differentiation, western blot quantification of σ^54^ and σ^28^-regulated proteins (Tarp [indirect] and HctB, respectively) showed reduced levels while the σ^66^-regulated late protein HctA levels were normal. Note that previous transcription experiments looking at *hctB* expression when altering PSM component levels did not show alterations (38, 39). Those experiments looked at earlier time points (12, 18, 24 and 30 hpi) than our study (48 hpi). TEM images of infected cells corroborated a developmental block in RsbV2 S55A expressing bacteria, which were predominantly in the RB or IB form compared to matched, uninduced samples that were enriched with EBs. The block in differentiation from IB to EB suggests that secondary differentiation has commenced in RsbV2 S55A conditions, likely through σ^66^-regulated late gene products such as HctA, but the production of σ^54^-and σ^28^-regulated late genes has not occurred to enable completion of secondary differentiation. Our results confirm the hierarchy of differentiation with σ^66^-regulated genes initiating secondary differentiation followed by σ^54^-regulated genes, and ultimately the σ^28^ regulon (46, 67).

With phosphorylation as the main driver of AASF function, regulation of RsbU and RsbW enzymatic activity becomes paramount for PSM output. These proteins appear to be regulated by the TCA intermediate α-ketoglutarate (RsbU, periplasmic sensor), the glycolytic intermediate PEP (RsbU, sensed by its cytoplasmic PP2C), and ATP (RsbW), supporting that nutrition/energy levels are critical for controlling PSM output (38–41). *Chlamydia* can obtain ATP from the host cell and via synthesis through substrate-level phosphorylation and a sodium dependent ATP synthase, which would allow the PSM to sense ATP levels of the host (indirectly) and the bacterium, while PEP would be an internal metabolite and α-ketoglutarate an external metabolite (68–73). Collectively, the PSM should be able to monitor energy levels of the host and bacterium to regulate bacterial growth and secondary differentiation through σ^66^ abundance. A variety of models have been proposed to explain the timing of secondary differentiation, including a nutrient-insensitive model (7, 74–77). Our data suggest that nutrient-levels should play a role in timing of differentiation, although we do not think that the PSM is the sole component dictating secondary differentiation events.

Absence of a phenotype when overexpressing RsbV1 S56A remains perplexing. Even with the reduced levels of RsbV1 S56A compared to RsbV2 S55A, RsbV1 S56A is a better binding partner for RsbW *in vitro*, and overexpressing RsbV2 S55A resulted in a very strong, deleterious phenotype (3 log reduction in IFUs). Interestingly, there is evidence in other bacteria that the phosphorylated AASF is involved in cell signaling and regulation of other factors beyond the ASF. In these cases, AASF knockout or expression of a non-phosphorylatable mutant did not result in a phenotype, similar to our findings with RsbV1. In *Sinorhizobium meliloti*, the phosphorylated AASF promotes the synthesis of mixed-linkage β-glucans through activation of a diguanylate cyclase by an unknown mechanism, and the non-traditional AASF from *Pseudomonas aeruginosa* interacts with FlgM in its non-phosphorylated form and c-di-GMP synthase in the phosphorylated form (11, 19, 78). There are additional examples of an AASF moonlighting outside of the PSM including regulation of the T3SS in *Bordetella* sp., hormogonia motility development in *Nostoc punctiforme*, biofilm formation in *Vibrio fischeri*, and production of a gene transfer agent (phage-like particle) and stationary phase viability in *Rhodobacter capsulatus* (20, 79–82). We hypothesize, similar to these other PSM systems, that RsbV1 has a moonlighting role and that its primary function may be to regulate an unknown target upon phosphorylation by RsbW. Such a scenario would leave RsbV2 as the canonical AASF within the chlamydial PSM. Attempts to identify non-PSM partners for RsbV1 and RsbV2 (plus/or minus phosphorylation) are ongoing. Collectively, the dual AASFs in the pathogenic *Chlamydia* have non-redundant functions with each playing roles that differ from the “ancestral” RsbV to help facilitate chlamydial endosymbiosis with new hosts.

## Material and Methods

### Bioinformatic Analyses

Protein sequences for PSM component homologs were obtained from the NCBI database (https://www.ncbi.nlm.nih.gov/, IDs listed in Table S2). AlphaFold predicted protein structures were downloaded from UniProt (https://www.uniprot.org/). Crystal protein structures of homologous proteins from *Geobacillus stearothermophilus* were obtained from the PDB database (https://www.rcsb.org/, SpoIIAB-σB [1L0O] and SpoIIAB-SpoIIAA [1TID] (30, 33). Structures were visualized using UCSF chimera (83), and the AlphaFold-derived chlamydial protein structures were superimposed with the crystal structure template followed by removal of the template. Amino acid sequences for AASF homologs from multiple species were aligned using Multiple Sequence Comparison by Log-Expectation (MUSCLE, https://www.ebi.ac.uk/Tools/msa/muscle/). The alignment was visualized in BioEdit version 7.2.5 using a 40% cut off for similarity/identity (84). The phylogenetic tree was constructed using Molecular Evolutionary Genetics Analysis (MEGA11) with maximum likelihood with 1000 bootstraps ((85), https://www.megasoftware.net/). Sequence similarities were mapped using CLuster ANalysis of Sequences (CLANS, (86), https://toolkit.tuebingen.mpg.de/tools/clans) using the BLOSUM62 scoring matrix and E-values of 1e^-4^. For the PSM protein map, proteins were connected using P<-1e^-4^, and P<-1e^-26^ was used for the for AASF protein map.

### Cell, chlamydial, and *E. coli* culture conditions

HeLa cells were routinely passaged in Dulbecco’s Modified Eagles Medium (DMEM) with 10% fetal bovine serum (FBS) at 37°C with 5% CO_2_. Cells were tested for mycoplasma contamination by PCR. A plasmid-free *C. trachomatis* L2 strain was used as the parent for recombinant strain construction (strains and vectors are listed in Table S3). Recombinant chlamydial strains were passaged in the presence of 500 µg/ml of spectinomycin for p2TK2-derived vectors (87) or 0.5 µg/ml of ampicillin for pBOMB-derived vectors (88), and titers were measured using the inclusion forming unit assay (IFU) using intrinsic fluorescence from the vector-derived GFP (pBOMB-series) or RFP (p2TK2-series) (89).

HeLa cells were infected with *C. trachomatis* at an MOI of 0.5 in infection medium (DMEM, 10% FBS, 1X non-essential amino acids, 400 ng/ml cycloheximide, and the appropriate antibiotic) for IFU and qPCR assays. An MOI of 0.25 was used for immunofluorescence assays with coverslips in 24 well plates. Infection was initiated through centrifugation at room temperature at 500xg for 30 min for 24 well plates or 60 min for 96 well plates. When needed, *rsbV1*/*rsbV2* gene expression was induced by adding 10 nM anhydrotetracycline (aTc) at 4 hpi except for experiments with C-terminal 3XFLAG-tagged RsbV1 S56A which used 112 nM aTc. For CRISPRi experiments, 2 nM of aTc was added at 4 hpi. To measure infectious progeny production from recombinant strains, 24 well plates were used. Well contents were harvested at 24 or 48 hpi by collection of supernatants followed by cell lysis via the addition of sterile water. Supernatant and lysis samples were pooled and bacteria pelleted by centrifugation at 4°C at 13,000xg for 5 min. Pellets were then suspended in 200 µl for RsbV2 (wild type and S55A) induction and 500 μl for other strains, and store at −80°C. To collect samples at multiple time points, 96 well plates were used as described for the 24 well plates. The lysed pellets were then suspended in 75 µl SPG for titering or 200 µl PBS for qPCR, and stored at −80°C. For western blot samples, cells were infected at an MOI of 0.5 and samples were harvested at 24, 36, and 48 hpi from 24 well plates using 150 µl of Laemmli buffer with β-mercaptoethanol to directly lyse the cells followed by heating at 95°C for 5 min.

*E. coli* strains DH5a, XL-1 Blue, or NEB10 were used use for cloning, while *E. coli* BL21(de3) or NEB express were used for protein production. Bacteria were grown in lysogeny broth (LB) or on LB agar plates at 37°C supplemented with antibiotics as needed: 100 µg/ml ampicillin, 50 µg/ml spectinomycin, and/or 34 µg/ml chloramphenicol.

### Quantification of *C. trachomatis* gDNA

Infected HeLa cells were processed as described in (41). Genomic DNA was isolated, and 50 ng of purified DNA was mixed with 0.3 µM of *omcB* forward and reverse primers with Power SYBR Green PCR Master Mix (Applied Biosystems). The qPCR reactions were run on a QuantStudio 3 (Applied Biosystems). The data were normalized using a standard curve from gDNA extracted from purified EBs.

### Vector construction

Genes were amplified by PCR using Phusion master mix (Thermo Scientific) (primers are listed in Table S4) or obtained as synthetic gBlocks (Integrated DNA Technologies, Table S5) and inserted into vectors using standard restriction cloning procedures, NEBuilder DNA HIFI assembly (New England BioLabs), or ligation independent cloning (pLATE31). The pLATE31 (Thermo Scientific) and pACYC^TM^-1 duet (Novagen) vectors were obtained from commercial sources. The *E. coli*-*C. trachomatis* shuttle vector p2TK2_Spec_-SW2 mCh(Gro_L2_) Tet-IncV-3xFLAG (kindly gifted from Isabelle Derré) was used as a template for chlamydial expression vector construction (87). For gene knockdown, the pBOMBL12CRia::L2 vector was used (57). For site directed mutagenesis, 5’-phosphorylated primers were used to amplify DNA followed by treatment with DpnI and self-ligation with T4 DNA ligase. All vectors were purified using the GeneJet Plasmid Kit (Thermo Scientific) and sequence verified by Sanger sequencing (PSOMAGEN). Vectors transformed into *C. trachomatis* were reisolated by transforming *E. coli* NEB10 with DNA isolated from recombinant strains (via the Whole Blood Genomic DNA Purification Kit, Thermo Scientific) followed by selection on LB agar with the appropriate drug. Plasmids were then isolated and checked for size on agarose gels, and the insert sequence reconfirmed by Sanger sequencing.

### Immunofluorescence microscopy

Cells were fixed at 24, 36, or 48 hrs (for 3XFLAG RsbV2/3xFLAG RsbV2 S55A induced with 10 nM aTc) or 48 hrs only (for 3XFLAG RsbV1 S56A/3XFLAG RsbV2 S56A induced with 112 nM aTc) in 3% paraformaldehyde and 3% sucrose in PBS and then permeabilized with 0.2% Triton X-100 in PBS. *Chlamydia* were detected using a mouse anti-MOMP primary antibody (1:250, Abcam ab41193) followed by staining with a donkey anti-mouse IgG-Alexa Fluor 488 secondary antibody (1:2000, Invitrogen A11001). FLAG-tagged proteins were detected with an anti-FLAG antibody (1:500, Sigma F1804). Samples were also stained with DAPI (300 nM) to detect DNA. Images were taken with a Leica DMi8 at 1000x with oil immersion and processed with ImageJ (NIH). Inclusion measurements were taken from at least two independent experiments and a minimum of 90 inclusions were counted in total.

### Protein detection via western blot

Laemmli-treated, *C. trachomatis* infected-cell samples were resolved on 15% SDS-PAGE gels for RsbV1, RsbV2, and HctA, 12% SDS-PAGE gels for HctB, or 6% SDS-PAGE gels for Tarp. Protein was then transferred to nitrocellulose membranes followed by blocking with 5% milk-Tris buffered saline (MTBS) for one hour at room temperature or overnight at 4°C (HctB blots). Primary antibodies diluted in MTBS were used to probe blots overnight at 4°C: polyclonal rabbit anti-RsbV2 antibody (1:200, (41)), polyclonal rabbit anti-RsbV1 antibody (1:200, (41)), monoclonal mouse anti-FLAG antibody (Sigma F1804, 1:1000), mouse anti-MOMP antibody (Abcam ab41193, 1:1000), rabbit anti-HctA antibody (kindly gifted by Ted Hackstadt, 1:1000), rabbit anti-HctB antibody (kindly gifted by Ted Hackstadt, 1:1000), or rabbit anti-Tarp antibody (kindly gifted by Ted Hackstadt, 1:100 000, (90)). Blots were then washed with 0.05% tween-20, Tris buffered saline (TTBS) and probed with the appropriate secondary antibody in MTBS; goat anti-mouse IgG conjugated with peroxidase (Sigma AP124P, 1:1000) or goat anti-rabbit IgG conjugated with peroxidase (Invitrogen 32260, 1:2000). Secondary-probed blots were washed three times with TTBS followed by TBS and incubated with Immobilon chemiluminescent substrate (Millipore). Images were acquired using a ChemiDoc Imager (Bio-Rad) and analyzed using Image Lab software (Bio-Rad). To quantify protein levels, the last image prior to band overexposure was used with MOMP levels serving as the loading control.

### Purification of recombinant proteins

Purification of recombinant GST-tagged RsbW and RsbU, or His-tagged RsbV1 and RsbV2 was performed as detailed in (41). The 6xHis-3xFLAG tagged proteins were purified using HisPur cobalt resin (Pierce) as previously described (41). Proteins were induced with 0.3 mM IPTG at 18°C for 20 h (His-3XFLAG-RsbV1) or with 0.3 mM IPTG at 30°C for 4 h (His-3XFLAG-RsbV2). Following purification, the 6xHis-tag was cleaved using the WELQUT protease (Thermo Scientific) in 100 mM Tris (pH 8.0) at 4°C overnight. The cleaved 6xHis-tag and WELQUT protease were removed using HisPur cobalt resin and the supernatants were retained. Preparation of GST-cleaved RsbU and phosphorylated RsbV1 and RsbV2 were performed as detailed in (41). Proteins were separated on SDS-PAGE gels to confirm purity and expected product sizes (see Supplemental Information and Figure S9). *In vitro* protein assays were performed with at least two independently purified protein preparations.

### *In vitro* kinase, phosphatase, and oligomerization assays

RsbW kinase and RsbU phosphatase assays were adapted from our previous studies (41). In brief, 2 µg or 3 µg of the substrate was mixed with enzyme at molar ratios of 1:0.19 (substrate:enzyme) for kinase assays or 1:0.4 (substrate:enzyme) for phosphatase assays. The kinase reaction was initiated by adding 1 mM ATP. Reactions were performed at 30°C and stopped by adding Laemmli buffer with β-mercaptoethanol and heating at 95°C for 5 min. The phosphorylation status of RsbV1/V2 was observed by running samples on SDS-PAGE gels (16% for His-V1/V2) supplemented with 20 µM of Phos binding reagent (APExBIO). Gels were stained with Coomassie brilliant blue and visualized with a Chemidoc Imager.

### TEM

HeLa cells were infected at an MOI of 1 with the recombinant *C. trachomatis* L2 strain possessing the plasmid encoding inducible expression of the FLAG-tagged RsbV2 S55A mutant. At 4 hpi, expression of the construct was induced or not with 10 nM aTc. At 48 hpi, samples were fixed with 2% glutaraldehyde, 2% paraformaldehyde in 0.1 M Sorensen’s phosphate buffer, pH 7.2. At the time of processing, samples were washed with PBS to remove excess fixative, and post fixation was performed for 1 h using a 1% aqueous solution of osmium tetroxide. Samples were dehydrated in a graded ethanol series (50%, 70%, 90%, 95%, and three times of 100% ethanol) with all steps being incubated for 15 min each. Subsequently, samples were soaked in 100% propylene oxide for 3 times for 15 min each. Samples were kept overnight in the fume hood while soaking in propylene oxide:Embed resin solution (1:1 ratio) until all the propylene oxide was evaporated. The following day, the samples were incubated in fresh resin for 2 h at room temperature, final embedding was performed, and samples polymerized overnight at 65°C. 90-nm thick sections of the polymerized blocks were cut on a Leica UC7 Ultramicrotome using a Diatome diamond knife. Sections were placed on uncoated 200 mesh copper grids, and staining was performed using 2% uranyl acetate followed by Reynold’s lead citrate. Sections were examined on a transmission electron microscope (Tecnai G2 Spirit TWIN FEI) operated at 80 kV, and images were acquired using an AMT digital imaging system.

### Statistics

GraphPad Prism 6.07 was used to perform all statistical analyses and p-values <0.05 were considered significant. Specific statistical tests and replicates are denoted in the respective figure legends.

## Acknowledgements

We thank Dr. José Vargas-Muñiz (Virginia Tech) for helpful discussion and Avigail A. Amezquita-Anaya (SIUC) for technical assistance. We also thank Dr. Ian Clarke (University of Southampton) for the plasmidless *C. trachomatis* L2 strain, Dr. Ted Hackstadt (NIH - Rocky Mountain Laboratories) for providing antibodies, and Dr. Isabelle Derré (University of Virginia) for providing vectors. The EMCF (Electron Microscopy Core) at UNMC is supported by state funds from the Nebraska Research Initiative (NRI) and the University of Nebraska Foundation and institutionally by the Office of the Vice Chancellor for Research. Research was supported by grant 2R15AI109566 (to D.J.F) and 1R21AI178150 (to S.P.O.) from the NIAID/National Institutes of Health. The content is solely the responsibility of the authors and does not necessarily represent the official views of the National Institutes of Health.

